# Higher Impulsivity in the Heterogeneous Structure of Theft Recidivists with and without Kleptomania

**DOI:** 10.64898/2026.07.27.740837

**Authors:** Yukiori Goto, Romane Charpentier, Satoshi Yoshino, Chikara Kita, Moojun Won, Young-A Lee

**Affiliations:** Graduate School of Informatics, Kyoto University, Kyoto, Japan; Faculty of Science, McGill University, Montreal, QC, Canada; Training Institute for Correctional Officials, Ministry of Justice of Japan, Tokyo, Japan; Shogakukan Academy Co., Ltd., Tokyo, Japan; MRC Lab Clinic, Tokyo, Japan; Department of Food Science and Nutrition, Daegu Catholic University, Gyeongsan, Korea

**Keywords:** Neurocriminology, Kleptomania, Behavioral addiction, Impulse control disorder, Psychiatric disorder, Impulsivity, Prefrontal cortex

## Abstract

Impulsivity is associated with various maladaptive behaviors, although its specific role in criminology has remained largely unexplored. We investigated impulsivity in theft recidivists (TR) with and without a diagnosis of kleptomania (KA) compared to control (CT) subjects with no criminal record in this study. Impulsivity was measured using a 5-trial adjusting delay discounting task, and prefrontal cortical (PFC) hemodynamics were assessed using functional near-infrared spectroscopy. Self-report questionnaires were also administered to further evaluate the relationships between impulsivity, negative affect, and reward and punishment sensitivity. TR demonstrated significantly higher impulsivity than CT, along with altered dorsomedial PFC hemodynamic responses. In addition, negative affect was significantly higher in the TR than in the CT participants. Path analysis revealed that the moderation of negative affect on reward, but not punishment, sensitivity, mediated impulsivity in TR. Notably, heightened impulsivity was observed across the TR participants regardless of KA diagnosis, whereas severe depression specifically distinguished KA from TR without it. These findings suggest that while trait impulsivity associated with PFC alterations may be a universal feature of TR, recurrent theft may be a heterogeneous condition, with specific affective dysregulations contributing differently to TR.

**SIGNIFICANCE STATEMENT:** Recurrent thefts, such as shoplifting, are a devastating social problem, posing massive financial losses. Nonetheless, surprisingly nothing is known about the neurobehavioral mechanisms that drive people stealing. In fact, kleptomania is a psychiatric disorder that had been identified over 200 years ago, yet exceptionally few studies have investigated this psychiatric condition, and its mechanism remains essentially unknown to date. This study demonstrates that impulsivity is elevated among incarcerated individuals with recurrent thefts with and without a diagnosis of kleptomania, which is associated with the activity of the left dorsomedial prefrontal cortex. Conversely, heightened depression among negative affect is a unique characteristic to kleptomania, providing evidence of heterogeneity among theft recidivists.

## 1. INTRODUCTION

Thefts and other property crimes are prevalent societal issues. Nevertheless, the neurobehavioral mechanisms underlying recurrent offending remain largely unknown. Thefts could also be related to kleptomania, a psychiatric condition with impulse-control disorder, characterized by recurrent failures to resist the stealing urges [1]. Although individuals with kleptomania frequently exhibit high comorbidity with mood disorders [2], the clinical and neurobiological boundaries separating kleptomania from other theft recidivism have not been well-defined.

Impulsivity has been suggested as a psychological risk factor for criminal offences [3–6]. Delay discounting paradigms, which measure the devaluation of future rewards as a function of temporal delay in favor of immediate, smaller rewards, are often used to assess impulsive decision-making, the tendency to engage in risky maladaptive behaviors without adequate consideration for future consequences [7,8]. Although such a delay discounting paradigm to examine impulsive decision-making has been administered in forensic populations [4], few studies have investigated impulsivity in the context of theft recidivism.

Impulsivity also interacts with emotional regulation. For instance, negative urgency, the tendency to engage in rash, reckless behaviors when experiencing distress, highlights the relationship between impulsivity and negative affect [9–11]. Studies have demonstrated that high negative urgency plays a role in psychiatric disorders, such as substance use disorders, so that impulsive actions alleviate negative emotional states in these individuals [9–11]. In addition, impulsivity could be associated with reward and punishment sensitivity [12–14]. Reward and punishment sensitivity are emotional drives that pursue rewards and evade negative consequences, respectively. As such, impulsive decision-making could be explained by augmented sensitivity to immediate rewards combined with dysregulated or diminished sensitivity to potential future punishments [12–14].

The prefrontal cortex (PFC) plays a major role in inhibiting impulsive and immediate urges [15,16]. Neuroimaging studies utilizing functional magnetic resonance imaging and near-infrared spectroscopy (fNIRS) have demonstrated that deficits in regions such as the dorsomedial and dorsolateral PFC are associated with behavioral disinhibition, poor decision-making, and increased risk-taking in neurological and psychiatric patients [17–19]. However, there remains a significant gap in neuroscientific research that specifically measures PFC responses to theft recidivism.

This study aimed to determine whether impulsivity is associated with recurrent theft, particularly shoplifting. We hypothesized that impulsivity was stronger in theft recidivists than in noncriminal controls. To evaluate this hypothesis, we investigated impulsivity and related factors (negative affect, reward and punishment sensitivity), alongside fNIRS measurements of PFC responses to unearth the neurobehavioral substrates of recurrent theft and kleptomania.

## 2. METHODS

### 2.1 Subjects

A total of 63 theft recidivists (TR) were recruited for this study. All TR subjects met the inclusion criteria of having prior records of incarceration due to larceny, primarily shoplifting, who were 18-79 years old and living in Japan at the time of the investigation. One participant dropped out of the entire investigation, and additional 5 participants dropped out of a part (ADT-5 task below) of the investigation, due to loss of interest in participation. Among the 62 TR subjects, 16 had been diagnosed with kleptomania (TR-KA) and were receiving treatment at clinics at the time of the investigation. As for a control (CT), 53 subjects who met the inclusion criteria of no criminal records at the age of 18-79 years old living in Japan were also recruited. None of the patients met the exclusion criteria for incompetence to understand the investigation details owing to intellectual disability or any other reasons. Age, sex, and smoking status were requested upon enrollment in the study. Age (Mann-Whitney U test, U = 1704, p = 0.736, Rank-Biserial Correlation r = -0.037) and sex (χ^2^ continuity correction, χ^2^(1) = 0.211, p = 0.646) did not differ between the TR and CT groups, whereas smoking was more prevalent in the TR group than in the CT group (χ^2^(1) = 15.23, p < 0.001). The sample structures are shown in Table 1.

**Table 1.** Sample structure regarding gender, age, smoking status, and Kleptomania Symptom Assessment Scale (K-SAS)

| Group | Gender | n | Age | Smoking* | K-SAS |  |  |  |  |
| --- | --- | --- | --- | --- | --- | --- | --- | --- | --- |
| | | | mean ± s.e.m.<br>(Min – Max) | n (%) | Total | Urge | Thought | Emotion | Stress |
| TR | Male | 29 | 43.2 ± 3.77<br>(18 – 76) | 23 (79.3) | 9.21 ± 2.05 | 2.21 ± 0.65 | 2.17 ± 0.61 | 2.17 ± 0.49 | 2.28 ± 0.50 |
| | Female | 33 | 51.4 ± 2.50<br>(18 – 74) | 9 (27.3) | 9.70 ± 1.21 | 2.09 ± 0.45 | 2.45 ± 0.38 | 2.64 ± 0.42 | 2.33 ± 0.42 |
| | Total | 62 | 47.5 ± 2.25<br>(18 – 76) | 32 (51.6) | 9.47 ± 1.15 | 2.15 ± 0.38 | 2.32 ± 0.35 | 2.42 ± 0.32 | 2.31 ± 0.32 |
| TR-KA | Male | 5 | 50.6 ± 5.30<br>(38 – 66) | 2 (40.0) | 9.00 ± 3.65 | 1.60 ± 0.81 | 3.00 ± 1.48 | 1.80 ± 1.11 | 2.00 ± 0.95 |
| | Female | 11 | 42.4 ± 4.46<br>(18 – 66) | 3 (27.3) | 9.45 ± 1.46 | 2.27 ± 0.56 | 2.91 ± 0.49 | 2.09 ± 0.65 | 2.00 ± 0.80 |
| | Total | 16 | 44.9 ± 3.53<br>(18 – 66) | 5 (31.3) | 9.31 ± 1.45 | 2.06 ± 0.45 | 2.94 ± 0.54 | 2.00 ± 0.55 | 2.00 ± 0.61 |
| TR-0 | Male | 10 | 52.0 ± 7.07<br>(20 – 76) | 10 (100.0) | 0 | 0 | 0 | 0 | 0 |
| | Female | 6 | 63.3 ± 2.96<br>(52 – 72) | 3 (50.0) | 0 | 0 | 0 | 0 | 0 |
| | Total | 16 | 56.4 ± 4.69<br>(20 – 76) | 13 (81.3) | 0 | 0 | 0 | 0 | 0 |
| CT | Male | 15 | 50.1 ± 3.81<br>(31 – 72) | 4 (26.7) | 0 | 0 | 0 | 0 | 0 |
| | Female | 38 | 48.2 ± 2.43<br>(20 – 78) | 4 (10.5) | 0 | 0 | 0 | 0 | 0 |
| | Total | 53 | 48.8 ± 2.03<br>(20 – 78) | 8 (15.1) | 0 | 0 | 0 | 0 | 0 |

K-SAS – Kleptomania symptom assessment scale; TR – Theft recidivist; TR-KA – Theft recidivist with kleptomania; TR-KSAS0 – Theft recidivist who scored 0 in K-SAS; CT – Control subjects with no criminal record
\*Subjects who were habitually smoking at the time and who used to have smoking habits in the past.

### 2.2 Questionnaire

Multiple self-report questionnaires were administered to assess stealing-related ideation, negative affect, and reward and punishment sensitivity. These questionnaires were in Likert format and translated into Japanese from the original English versions by the author (YG) at the time of the current study or had been translated and used in our previous study [20].

#### 2.2.1 Kleptomania Symptom Assessment Scale (K-SAS)

Stealing ideation was evaluated using K-SAS [21], which is the 11-item scale to quantify a state of kleptomania symptom severity over the past one week. The scale consists of four subscales to evaluate different dimensions: urge and impulses, thoughts, emotional and psychological response, and distress and impairment. The K-SAS scores ranged from a minimum of 0 to a maximum of 44, along with 8-20 as mild, 21-30 as moderate, and 31-40 as severe symptom levels, respectively. In this study, the K-SAS was used for all participants, regardless of whether they had been diagnosed with KA. The K-SAS was also administered to the CT group to confirm that all participants in this group scored 0 on the scale.

#### 2.2.2 Depression, Anxiety, and Stress Scale, 21 item version (DASS-21)

Negative affect was evaluated using the DASS-21 [22]. The questionnaire was designed to measure states of negative affect over the past week, with a set of three domains: stress, anxiety, and depression. Overall negative affect was expressed by combining the scores of these three subscales.

#### 2.2.3 Reward and Punishment Responsivity and Motivation Questionnaire (RPRM-Q)

Reward and punishment sensitivity was assessed using the RPRM-Q [23]. The questionnaire evaluates four dimensions of reward- and punishment-related behaviors: reward responsivity, reward motivation, punishment responsivity, and punishment motivation. Responsivity is defined as an emotional reaction to reward or punishment stimuli, whereas motivation is an emotional drive to actively pursue rewards or to evade punishments. Reward and punishment sensitivities are represented by combining the responsivity and motivation subscales.

### 2.3 5-Trial Adjusting Delay Discounting Task (ADT-5)

The ADT-5 [8] is a brief delay discounting task used to measure impulsive tendencies in decision making. In the task, participants were asked to choose between an immediate small reward and a delayed large reward in a series of five questions, with a variable delay for the large reward in each question. A discount rate k, in which a higher value indicates a higher impulsive tendency, was calculated based on the hyperbolic decay formula, V = A / (1 + kD), where V and A are the subject and actual values of the reward, respectively, and D is the delay until the large reward is received. Because the k-values had a heavily skewed distribution, their log transformation, log(k), was used for subsequent data analysis. The task was programmed and administered using the Inquisit Web (Millisecond Software, LLC).

### 2.4 fNIRS

PFC responses while the participants performed the ADT-5 task were measured using fNIRS. The NIRO-200 NIRS Image Processing and Measuring System (Hamamatsu Photonics K.K.) was used for the measurements. The system consisted of two emitters (E1, E2; delivering laser pulse at the wavelength of 775, 810, and 850 nm) and eight detectors (CH1‒CH8), with the distance at 3.0 cm apart between an emitter and a detector, with which oxygenated (O2Hb) and deoxygenated (HHb) hemoglobin changes over time with a sampling rate of 0.5 Hz at 10 locations spanning over the left and right hemisphere of PFC R1‒R10, where R4 and R6 covered the left and right rostral dorsal PFC (PFrd) corresponding to Brodmann area (BA) 10/9/8, R5 and R7 covered the left and right caudal dorsomedial PFC (PFcdm) corresponding to BA6/8, R3 and R8 covered the left and right rostral dorsolateral superior PFC (PFrdls) corresponding to BA10/9, R2 and R9 covered the left and right caudal dorsolateral PFC (PFcdl) corresponding to BA45/46/9, and R1 and R10 covered the intermediate zone between PFrdls and PFcdl, of the MarsAtlas [24].

### 2.5 Experimental Design

This study was conducted in accordance with the Declaration of Helsinki and the Ethical Guidelines for Medical and Health Research Involving Human Subjects by Japanese Ministry of Health, Labour and Welfare. All procedures were approved by the Human Research Ethics Committee of Kyoto University Graduate School of Informatics. Written informed consent was obtained from all participants before the investigations. Then, questionnaires were administered, followed by the ADT-5 task using fNIRS. fNIRS was conducted in all CT subjects but only in 39 TR subjects without the diagnosis of kleptomania due to environmental and logistical constraints.

### 2.6 Data Analysis

All data are expressed as mean ± standard error of the mean (s.e.m.). Statistical significance was set at p < 0.05. Statistical analyses were conducted using JASP ver. 0.97.1 [25] and OriginPro ver. 2026 (OriginLab Corporation, Northampton, MA, USA).

#### 2.6.1 Analysis of covariance (ANCOVA)

A comparison of dependent measurements for impulsivity, negative affect, and reward/punishment sensitivity between the groups (TR vs. CT) was conducted using ANCOVA, along with adjustment for age, sex, and smoking status of subjects as covariates. The interactions of the covariates with the group were examined using a process model. When dependent variables were found to violate normality (Shapiro-Wilk test for normality) and homogeneity (Levene’s test for equality of variance) by assumption checks, transformation of the data using Rank-Based Inverse Normal Transformation was conducted prior to statistical analysis. In addition, a post hoc nonparametric Kruskal-Wallis test was performed to compare the groups.

#### 2.6.2 Process model

Relationships for group differences in impulsivity with reward/punishment sensitivity and negative affect were evaluated using a process model. In this model, reward/punishment sensitivity was specifically tested as a mediator (stronger and weaker reward punishment/reward sensitivity may in turn facilitate impulsivity) and negative affect as a moderator (such as in the case of negative urgency).

##### Nonparametric statistical comparison

Comparisons of subgroups within TR were conducted using the nonparametric Mann-Whitney U test.

#### 2.6.3 Linear mixed-effect model (LMM)

LMM was used to assess group differences (TR vs. CT) in PFC responses measured with fNIRS to account for the non-independence of the data inherent in the repeated-measures design, where recordings over time were nested within each subject. Accordingly, the model is described by the following equation:

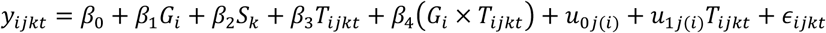

where *y_it_* is the observed outcome for Subject j (j = 1, 2,…) in Group i (i = 1, 2), with Signal k (k = 1, 2) at Time t, *G_i_* is the Group indicator (1 for CT, 2 for TR), *S_k_* is the Signal indicator (1 for O2Hb, 2 for HHb), *T_ijkt_* and is the continuous time point of the sampling. As for fixed effects, *β*_0_ is the overall fixed intercept (grand mean at baseline), *β*_1_ is the main effect of Group (the average difference between groups across all times), *β*_2_ is the main effect of Signal (the constant difference between oxy-Hb and deoxy-Hb, regardless of Group and Time), *β*_3_ is the main effect of Time (the average change over time across both groups), and *β*_4_ is the Group × Time interaction (the difference in the slope/trajectory between the groups). For random effects, *u_0j(i)_* is the random intercept for Subject j nested in Group i, *u*_1*j(i)*_*T_ijkt_* is the random slope for Time for Subject j, and *ε_it_* is the residual error. The Group × Time interaction term was the primary effect of interest for determining whether the trajectories of the outcomes over time differed significantly between the groups. To account for baseline variability among individuals, a random intercept for the subjects was included in the model. Models were fit using restricted maximum likelihood, and p-values for fixed effects were estimated using the Satterthwaite approximation. The p-values for the comparisons of the estimated marginal means between groups were adjusted using the Bonferroni method.

## 3. RESULTS

### 3.1 TR vs. CT

All CT subjects scored 0 on the K-SAS, whereas the K-SAS score in TR subjects was 9.47 ± 1.15 (mean ± s.e.m.; Wilcoxon signed-rank test to evaluate the score higher than 0, W = 1081, p < 0.001, r =1.00 ± 0.17; Table 1).

Comparisons of impulsivity, reward/punish sensitivity, and negative affect were conducted using ANCOVA, along with adjustments for age, sex, and the smoking status. The log of the discount rate k, log(k), in ADT-5 was significantly smaller (more negative) in TR than in CT, suggesting that TR participants made more impulsive choices (F(1, 110) = 13.47, p < 0.001, ω^2^ = 0.104; Suppl. Table S1; Fig. 1a). The group difference was further confirmed with Kruskal-Wallis test (Suppl. Table S1). The total (combining all stress, anxiety, and depression subscales) score of the DASS-21 was also significantly higher in the TR group than in bn the CT group, suggesting that TR participants had more severe negative affect at the time of investigation (F(1, 110) = 41.72, p < 0.001, ω^2^ = 0.262; Fig. 1b; Suppl. Table S1). Although the reward sensitivity (combining response and motivation subscales) score assessed with the RPRM-Q was not different between groups (Fig. 1c; Suppl. Table S1), the punishment sensitivity score was small (ω^2^ = 0.038) but higher in TR than in CT (F(1, 110) = 5.504, p = 0.021; Fig. 1d; Suppl. Table S1).

**Figure 1.**
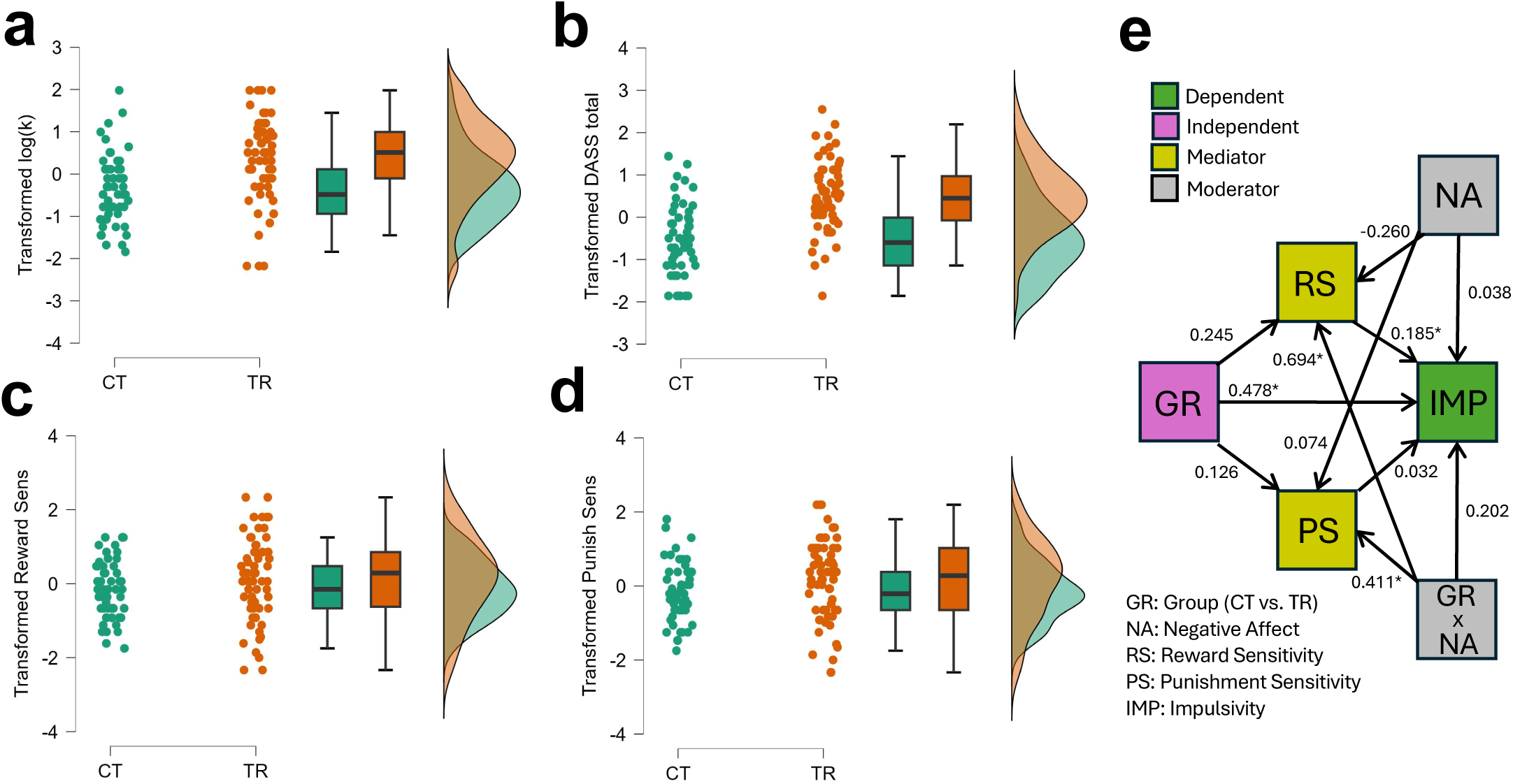
Comparisons of behavioral measurements between TR and CT. a-d, Raincloud plots comparing transformed (data with rank-based inverse normal transformation) impulsivity (a), negative affect (b), reward sensitivity (c), and punishment sensitivity (d) between TR and CT. **e,** Statistical path plot of the Process Model, with path coefficients.

To understand the underlying process associated with higher impulsivity in TR, the relationships between impulsivity, negative affect, and reward/punishment sensitivity were evaluated using a process-model. The model comprised the direct effect of the group on impulsivity, along with the indirect effects of reward/punishment sensitivity on impulsive behavior. Negative affect is also known to affect impulsivity, which could be through the moderation of both direct and indirect effects (Fig. 1e). The variances explained by the predictors in this model were R^2^ = 0.204 for impulsivity, R^2^ = 0.118 for reward sensitivity, and R^2^ = 0.135 for punishment sensitivity. A significant path coefficient was observed for the direct group effect on impulsivity (coefficient b = 0.478, Z = 2.433, p =0.015; Fig. 1e; Suppl. Table S2). The path coefficient for reward (b = 0.185, Z = 2.104, p = 0.035; Fig. 1e; Suppl. Table S2), but not punishment, sensitivity to impulsivity was also significant. Moreover, significant path coefficients were found for the interaction between group and negative affect to reward (b = 0.694, Z = 3.237, p = 0.001; Fig. 1e; Suppl. Table S2) and punishment (b = 0.411, Z = 1.970, p = 0.049; Fig. 1e; Suppl. Table S2) sensitivity, and thereby, a negative affect-dependent indirect group effect on impulsivity through reward sensitivity emerged (Fig. 1e; Suppl. Table S2). Accordingly, in probe conditional continuous effect assessments, this indirect effect of negative affect to impulsivity through reward sensitivity became more significant from participants with low to high levels of negative affect (16th percentile, which could be primarily CT participants; b = -0.082, Z = -1.230, p=0.219; 50th percentile, which could be a mixture of CT and TR participants; b = 0.043, Z = 0.983, p=0.326; 86th percentile, which could be primarily TR participants; b = 0.170, Z = 1.727, p=0.084; Suppl. Table S2). The direct group effect on impulsivity was also negative affect-dependent, such that a significant effect was found in the 50th (b = 0.475, Z = 2.421, p=0.015; Suppl. Table S2) and 86th (b = 0.674, Z = 2.284, p=0.022; Suppl. Table S2), but not 16, percentiles of negative affect severity (Suppl. Table S2).

Collectively, these results suggest that impulsivity may be higher in TR, which may be associated with the direct effect of negative affect on impulsivity and indirectly through its modulation of reward sensitivity.

### 3.2 TR-KA vs. TR-NK

Sixteen of the 62 TR participants were diagnosed with kleptomania (TR-KA). To determine the differences between those with and without kleptomania, comparisons between TR-KA and TR-NK (TR participants without a diagnosis of kleptomania) were conducted.

No group differences were observed in terms of age, sex, or smoking status (Table 1). Moreover, the K-SAS scores, including those for each subscale, did not differ between the groups (Table 1). The measurements for impulsivity, as well as reward and punishment sensitivity, were also not different between the groups (Suppl. Table S3).

However, although overall negative affect (total score of DASS-21) was not different between the groups, a score for the depression subscale, but not other stress and anxiety subscales, was higher in TR-KA than that of TR-NK (U = 503.5, p=0.029, r = -0.368; Suppl. Table S3).

These results suggest that higher impulsivity may be a trait of TR, regardless of whether KA is present, whereas higher depression may be a unique characteristic associated with KA.

### 3.3 Subgroup of TR

Among the 62 TR subjects, 16 rated 0 on the K-SAS (25.8%; n=2 out of 16 TR-KA, 12.5%; n=14 out of 46 TR-NK, 30.4%; Table 1). These TR subjects who scored 0 in K-SAS (TR-0) were significantly different from not only other TR subjects (TR-X, who scored higher than 0 in K-SAS) but also CT subjects (who also rated 0 in K-SAS) in their age (significantly older, U = 513.5, p = 0.020, r = -0.395, vs. TR-X; U = 552.5, p = 0.069, r = -0.303, vs. CT), sex (consisting of more males, χ^2^(1) = 7.26, p = 0.007, vs. TR-X; χ^2^(1) = 6.22, p = 0.013, vs. CT), and smoking (consisting of more smokers, χ^2^(1) = 6.07, p =0.014, vs. TR-X; χ^2^(1) = 25.4, p < 0.001, vs. CT), suggesting a possibility that these subjects may consist of a subgroup within RT.

Although impulsivity did not differ between TR-0 and TR-X (Suppl. Table S3), negative affect in TR-0 was significantly milder than that of TR-X, but still higher than that of CT (total score of DASS-21, U = 218.0, p = 0.016, r = 0.408, vs. TR-X; U = 578.5, p = 0.028, r = -0.364, vs. CT; Suppl. Table S3).

These results suggest that there may be a subgroup within TR that consists primarily of older male participants with a high smoking prevalence, who consider themselves to have no theft ideation and lower negative affect.

### 3.4 Impulsivity-related PFC Responses

fNIRS was used to measure PFC responses in 53 CT and 39 TR (only those without kleptomania) participants while they engaged in the ADT-5 task.

LMM was conducted with groups (CT vs. TR), signals (O2Hb vs. HHb), and time (sampling over time) as fixed effects and subjects as random effects. The LMM was performed independently for each sampling site R1-R10. A significant group × time interaction was found at R4, corresponding to the left rostrodorsal PFC (F(1, 83.74) = 5.998, p = 0.016, Vovk-Sellke maximum p-ratio (VS-MPR) = 5.455; Fig. 2; Suppl. Table S5). Significant changes in signals over time were also observed in R5, corresponding to the left caudal dorsomedial PFC, the region anterior to R4, although no group or group × time difference was found in this area (Suppl. Table S5).

**Figure 2.**
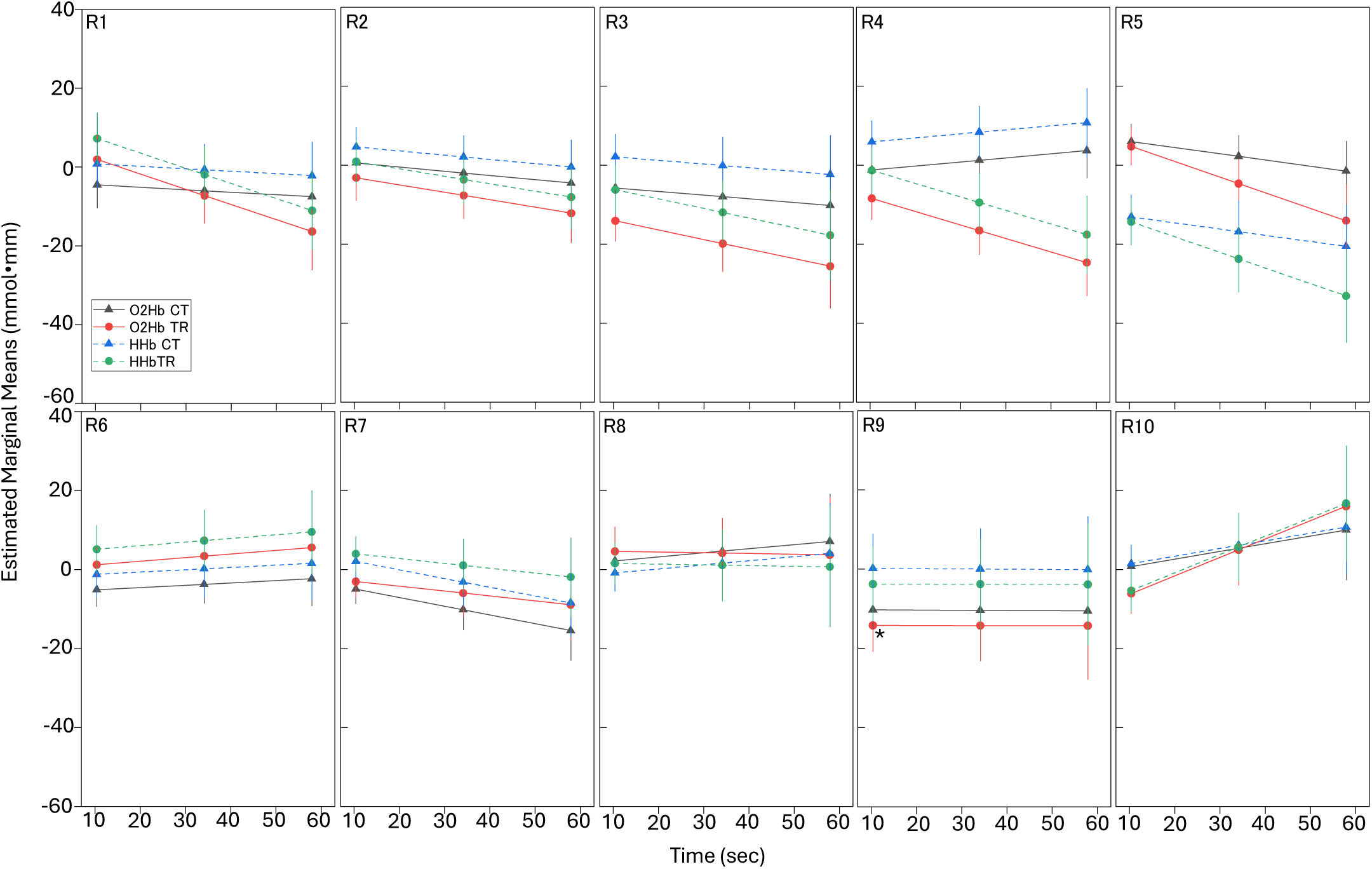
Marginal estimated means of fNIRS signals with linear mixed-effects model. Panels showing the marginal mean estimates of oxygenated hemoglobin (O2Hb) and deoxygenated hemoglobin (HHb) signal changes over time at the sampling sites R1-R10. Error bars indicate s.e.m.

Collectively, these results suggest that the left dorsomedial PFC, particularly rostrodorsal region, may be associated with higher impulsive decision-making in the TR group.

## 4. DISCUSSION

This study investigated the neurobehavioral mechanisms underlying recurrent theft, specifically focused on impulsivity. Behavioral assessments using the delay discounting task revealed that theft recidivists exhibited significantly higher impulsivity, alongside elevated levels of negative affect, compared to control subjects with no criminal records. With statistical path analysis, we further clarified that in theft recidivists, negative affect indirectly accentuates impulsive decision-making through the negative affect-dependent modulation of reward sensitivity. In particular, although augmented impulsivity remained ubiquitous across all theft recidivists, severe depression emerged as a unique and distinguishing state specifically observed in theft recidivists diagnosed with kleptomania. We further identified psychological heterogeneity within theft recidivists, isolating a distinct demographic subgroup characterized by the absence of subjective stealing ideations, which was typical of older males with a high prevalence of smoking and milder negative affective states. fNIRS identified a significant group-by-time hemodynamic interaction localized in the left dorsomedial PFC during impulsive choices, which is partly consistent with other studies demonstrating the role of this area in controlling impulsive behaviors [17–19].

Our findings align with and expand upon the neurobehavioral mechanisms suggested for maladaptive behaviors among criminal offenders. The identified relationship, wherein negative affect amplifies impulsivity both directly and via reward-system interactions, supports the theoretical framework of negative urgency, which posits that individuals with poor impulse control frequently engage in rash, risky actions as a maladaptive strategy to rapidly escape intense emotional distress [9–11]. Moreover, the finding that severe depressive states uniquely characterize individuals with KA mirrors clinical studies highlighting the frequent comorbidity between KA and major depressive disorder [2]. However, our findings differ partially from the mechanisms suggested for other forensic subjects with psychiatric conditions, such as addictive disorders. While typical addictive disorder frameworks assume hypersensitivity to rewards and deficits in punishment processing [26–28], theft recidivists in this study demonstrated no deviations in reward sensitivity compared to control individuals and even showed a slight increase in punishment sensitivity.

Our findings further suggest that alongside KA, a distinct subgroup may also exist among theft recidivists without KA. Individuals in this subgroup demonstrated an absence of stealing ideation as measured by the K-SAS; however, this must be interpreted with caution given the subjective assessment nature of the scale. Notably, this subgroup exhibited advanced age and an exceptionally high prevalence of smoking. These observations indicate that their behavioral presentations may closely mirror the profiles of individuals with substance use disorders or dementia, particularly frontotemporal dementia, in which antisocial behaviors such as recurrent stealing often manifest [29].

This study had several limitations. First, the sample size was limited to a relatively small number, especially given the difficulty of recruiting subjects with the unique characteristic nature of theft and recidivism. Moreover, due to unavoidable environmental and logistical constraints during data collection, fNIRS could not be completed for the subgroup of theft recidivists diagnosed with kleptomania, limiting the interpretation of PFC hemodynamic comparisons associated with impulsivity in noncriminal control participants. Consequently, whether PFC deficits differ between compulsive and noncompulsive theft remains to be addressed. Second, the study relied on self-report scales to quantify negative affect, reward/punishment sensitivity, and recent stealing ideations, which may have introduced subjective reporting biases or strategic answers. In fact, we found that there was a subgroup of theft recidivists characterized by the absence of subjective stealing ideations with the K-SAS. Such heterogeneity is likely to be revealed as a consequence of subjective reports. Finally, due to the cross-sectional nature of the study design, it remains unclear whether higher impulsivity associated with PFC alterations and heightened negative urgency are pre-existing vulnerabilities that directly cause theft or recidivism. These findings may be a consequence of chronic legal distress, incarceration, and lifestyle instability.

In conclusion, our study provides novel insights into the field as the first direct neuroscientific evidence demonstrating that the neurobehavioral processes of recurrent theft are not monolithic. Thus, although trait impulsivity is higher regardless of a diagnosis of kleptomania, affective presentation is divergent. This study challenges the assumption that property offenders are uniform, thereby expanding our understanding of forensic typologies. In particular, fNIRS measurements of the PFC yielded an insight that functional abnormalities specifically within the left dorsomedial PFC may be associated with defective inhibition of impulsive thoughts and poor consequence evaluation, which could cause repetitive property offending, offering a potential biological target for intervention.

## List of Abbreviations

ADT-5: 5-Trial Adjusting Delay Discounting Task
ANCOVA: Analysis of Covariance
BA: Brodmann Area
CT: Control
DASS-21: Depression, Anxiety, and Stress Scale, 21 item version
fNIRS: Functional Near-Infrared Spectroscopy
HHb: Deoxygenated Hemoglobin
KA: Kleptomania
K-SAS: Kleptomania Symptom Assessment Scale
LMM: Linear Mixed-Effect Model
O2Hb: Oxygenated Hemoglobin
PFC: Prefrontal Cortex
PFcdl: Caudal Dorsolateral PFC
PFcdm: Caudal Dorsomedial PFC
PFrd: Rostral Dorsal PFC
PFrdls: Rostral Dorsolateral Superior PFC
TR: Theft Recidivists
TR-KA: Theft Recidivists with Kleptomania
TR-NK: Theft Recidivists without Kleptomania
TR-0: Theft Recidivists who rated 0 in K-SAS
TR-X: Theft Recidivists who rated higher than 0 in K-SAS
RPRM-Q: Reward and Punishment Responsivity and Motivation Questionnaire
VS-MPR: Vovk-Sellke Maximum P-Ratio

## Data Availability

Data is available from the corresponding author (YG) upon a reasonable request.

## Acknowledgements

We would like to thank the staff of the Non-Profit Organization Kurashi O-en Network (Nagoya, Japan), Nishi Hongwanji Byakkoso (Kyoto, Japan), Non-Profit Organization Kyoto MAC (Kyoto, Japan), Kyoto Hogo Ikusei Kai (Kyoto, Japan), and Liberty Women’s House Olive (Otsu, Japan) for recruiting participants, scheduling surveys, and providing various technical supports.

## Author Contributions

YG contributed to the study design, data collection, and analysis. RC contributed to the study design and analysis. SY and CK contributed to the study design and data collection. YAL contributed to the analysis. All authors have reviewed, edited, and approved the manuscript prior to submission.

## Funding

This work was supported by the Japan Society for the Promotion of Science (JSPS) Grant-in-Aid for Scientific Research (B) 25K00898, awarded to YG.

## Competing Interests

The authors declare no conflicts of interest.

## Notes

### Competing Interest Statement

The authors have declared no competing interest.

