## Supplementary Table S1-S5 for "Higher Impulsivity in the Heterogeneous Structure of Theft Recidivists with and without Kleptomania"

**Supplementary Table S1. A Summary Table of Statistical Comparisons between TR and CT with ANCOVA and Kruskal-Wallis Test for ADT-5, DASS-21, and RPRM-Q**

| | | mean ± s.e.m. | | ANCOVA | | | Kruskal-Wallis Test | | |
|---|---|---|---|---|---|---|---|---|---|
| | | TR | CT | $F_{(1, 110)}$ | p | Effect Size ($\omega^2$) | H | p | Rank Effect Size ($\varepsilon^2$) |
| **ADT-5** | log(k) | -2.93 ± 0.43 | -5.14 ± 0.34 | 13.47 | <0.001 | 0.104 | 17.32 | <0.001 | 0.159 |
| **DASS-21** | Total | 23.1 ± 1.42 | 10.3 ± 1.35 | 41.72 | <0.001 | 0.262 | 31.96 | <0.001 | 0.280 |
| | Stress | 9.26 ± 0.55 | 4.87 ± 0.59 | 26.42 | <0.001 | 0.180 | 23.01 | <0.001 | 0.202 |
| | Anxiety | 5.73 ± 0.52 | 1.94 ± 0.35 | 34.85 | <0.001 | 0.232 | 32.28 | <0.001 | 0.283 |
| | Depression | 7.92 ± 0.70 | 3.47 ± 0.54 | 29.60 | <0.001 | 0.196 | 19.57 | <0.001 | 0.172 |
| **RPRM-Q** | RR | 11.8 ± 0.47 | 12.0 ± 0.29 | 0.123 | 0.726 | 0.000 | 0.454 | 0.500 | 0.004 |
| | RM | 13.2 ± 0.56 | 11.9 ± 0.37 | 3.404 | 0.068 | 0.019 | 6.529 | 0.011 | 0.057 |
| | RS | 25.0 ± 0.95 | 23.9 ± 0.59 | 1.838 | 0.178 | 0.007 | 3.568 | 0.059 | 0.031 |
| | PR | 14.1 ± 0.69 | 13.2 ± 0.56 | 5.187 | 0.025 | 0.035 | 3.503 | 0.061 | 0.031 |
| | PM | 10.9 ± 0.44 | 9.77 ± 0.39 | 5.315 | 0.023 | 0.036 | 4.756 | 0.029 | 0.042 |
| | PS | 24.8 ± 1.08 | 22.9 ± 0.81 | 5.504 | 0.021 | 0.038 | 4.644 | 0.031 | 0.041 |

ADT-5 – 5-trial adjusting delay discounting task; DASS-21 – 21 item version of the depression anxiety stress scale; RPRM-Q – Reward and punishment responsivity and motivation questionnaire; TR – Theft recidivist; CT – Control subjects with no criminal record; RR – Reward response; RM – Reward motivation; RS – Reward sensitivity; PR – Punishment response; PM – Punishment motivation; PS – Punishment sensitivity

**Supplementary Table S2. A Summary Table of Statistics in the Process Model for the Relationships with Impulsivity, Negative Affect, and Reward/Punishment Sensitivity**

| | | | | | | NA | Estimate | Z | p |
|---|---|---|---|---|---|---|---|---|---|
| | | | | | | Percentile | mean ± s.e.m. | | |
| **Path Coefficients** | GP | → | IMP | | | | 0.478 ± 0.197 | 2.433 | 0.015 |
| | RS | → | IMP | | | | 0.185 ± 0.088 | 2.104 | 0.035 |
| | PS | → | IMP | | | | 0.032 ± 0.090 | 0.352 | 0.725 |
| | NA | → | IMP | | | | 0.038 ± 0.146 | 0.261 | 0.794 |
| | GP x NA | → | IMP | | | | 0.202 ± 0.211 | 0.959 | 0.338 |
| | GP | → | RS | | | | 0.245 ± 0.211 | 1.160 | 0.246 |
| | NA | → | RS | | | | -0.260 ± 0.156 | -1.668 | 0.095 |
| | GP x NA | → | RS | | | | 0.694 ± 0.215 | 3.237 | 0.001 |
| | GP | → | PS | | | | 0.126 ± 0.206 | 0.611 | 0.541 |
| | NA | → | PS | | | | 0.074 ± 0.152 | 0.489 | 0.625 |
| | GP x NA | → | PS | | | | 0.411 ± 0.209 | 1.970 | 0.049 |
| **Direct/Indirect Effects** | GP | → | IMP | | | 16 | 0.278 ± 0.274 | 1.014 | 0.311 |
| | GP | → | IMP | | | 50 | 0.475 ± 0.196 | 2.421 | 0.015 |
| | GP | → | IMP | | | 84 | 0.674 ± 0.295 | 2.284 | 0.022 |
| | GP | → | RS | → | IMP | 16 | -0.082 ± 0.067 | -1.230 | 0.219 |
| | GP | → | RS | → | IMP | 50 | 0.043 ± 0.044 | 0.983 | 0.326 |
| | GP | → | RS | → | IMP | 84 | 0.170 ± 0.098 | 1.727 | 0.084 |
| | GP | → | PS | → | IMP | 16 | -0.009 ± 0.027 | -0.331 | 0.740 |
| | GP | → | PS | → | IMP | 50 | 0.004 ± 0.013 | 0.301 | 0.763 |
| | GP | → | PS | → | IMP | 84 | 0.017 ± 0.048 | 0.345 | 0.730 |

GP – Group (TR vs. CT); NA – Negative affect; GP x NA – Group and negative affect interaction; RS – Reward sensitivity; PS – Punishment sensitivity; IMP – Impulsivity

**Supplementary Table S3. A Summary Table of Statistical Comparisons between TR-KA and TR-NK with Man-Whiteny U test for ADT-5, DASS-21, and RPRM-Q**

| | | mean ± s.e.m. | | U | p | Effect Size (r) |
|---|---|---|---|---|---|---|
| | | TR-KA | TR-NK | | | |
| **ADT-5** | log(k) | -3.18 ± 2.69 | -2.85 ± 3.38 | 277.5 | 0.669 | 0.078 |
| **DASS-21** | Total | 23.8 ± 2.82 | 22.8 ± 1.66 | 378.0 | 0.878 | 0.027 |
| | Stress | 8.19 ± 0.98 | 9.39 ± 0.69 | 311.5 | 0.366 | 0.154 |
| | Anxiety | 4.75 ± 1.05 | 6.76 ± 0.72 | 273.5 | 0.129 | 0.257 |
| | Depression | 10.2 ± 1.40 | 6.67 ± 0.67 | 503.5 | 0.029 | -0.368 |
| **RPRM-Q** | RR | 11.6 ± 0.70 | 11.9 ± 0.59 | 300.5 | 0.278 | 0.183 |
| | RM | 13.6 ± 0.80 | 13.1 ± 0.70 | 363.0 | 0.942 | 0.014 |
| | RS | 25.1 ± 1.29 | 24.9 ± 1.21 | 352.5 | 0.808 | 0.042 |
| | PR | 14.6 ± 0.89 | 13.9 ± 0.89 | 408.5 | 0.516 | -0.110 |
| | PM | 11.6 ± 0.38 | 10.6 ± 0.58 | 338.5 | 0.640 | 0.080 |
| | PS | 25.6 ± 1.38 | 24.5 ± 1.38 | 368.0 | 1.000 | 0.000 |

ADT-5 – 5-trial adjusting delay discounting task; DASS-21 – 21 item version of the depression anxiety stress scale; RPRM-Q – Reward and punishment responsivity and motivation questionnaire; RR – Reward response; RM – Reward motivation; RS – Reward sensitivity; PR – Punishment response; PM – Punishment motivation; PS – Punishment sensitivity; TR-KA – Theft recidivist with the diagnosis of kleptomania; TR-NK – Theft recidivist without the diagnosis of kleptomania; CT – Control subjects with no criminal record; r – Rank biserial correlation

**Supplementary Table S4. A Summary Table of Statistical Comparisons between TR-0 and other TR-X with Man-Whiteny U test for ADT-5, DASS-21, and RPRM-Q**

| | | mean ± s.e.m. | | U | p | Effect Size (r) |
|---|---|---|---|---|---|---|
| | | TR-0 | TR-X | | | |
| **ADT-5** | log(k) | -2.80 ± 0.90 | -2.98 ± 0.49 | 345.5 | 0.586 | -0.097 |
| **DASS-21** | Total | 16.8 ± 2.75 | 25.3 ± 1.55 | 218.0 | 0.016 | 0.408 |
| | Stress | 6.31 ± 1.05 | 10.0 ± 0.62 | 204.5 | 0.008 | 0.444 |
| | Anxiety | 5.50 ± 1.08 | 6.50 ± 0.73 | 327.5 | 0.518 | 0.110 |
| | Depression | 4.94 ± 1.04 | 8.50 ± 0.74 | 212.5 | 0.012 | 0.423 |
| **RPRM-Q** | RR | 11.1 ± 1.25 | 12.0 ± 0.47 | 358.0 | 0.878 | 0.027 |
| | RM | 11.9 ± 1.52 | 13.6 ± 0.53 | 322.5 | 0.467 | 0.124 |
| | RS | 23.6 ± 2.67 | 25.7 ± 0.89 | 230.5 | 0.027 | 0.374 |
| | PR | 11.3 ± 1.61 | 15.1 ± 0.71 | 297.0 | 0.252 | 0.193 |
| | PM | 9.44 ± 1.22 | 11.4 ± 0.40 | 340.5 | 0.663 | 0.075 |
| | PS | 20.8 ± 2.77 | 26.2 ± 1.03 | 262.0 | 0.089 | 0.288 |

ADT-5 – 5-trial adjusting delay discounting task; DASS-21 – 21 item version of the depression anxiety stress scale; RPRM-Q – Reward and punishment responsivity and motivation questionnaire; RR – Reward response; RM – Reward motivation; RS – Reward sensitivity; PR – Punishment response; PM – Punishment motivation; PS – Punishment sensitivity; TR-0 – Theft recidivist who scored 0 in K-SAS; TR-X – Theft recidivist who scored 1 or higher in K-SAS; CT – Control subjects with no criminal record; r – Rank biserial correlation

**Supplementary Table S5. A Summary of Statistical Results with Liner Mixed-effects Model (LMM) for Prefrontal Cortical (PFC) Responses Comparing TR and CT**

| | Group | | | Time | | | Group x Time | | | Signal (O2Hb vs. HHb) | | |
|---|---|---|---|---|---|---|---|---|---|---|---|---|
| | F | p | VS-MPR | F | p | VS-MPR | F | p | VS-MPR | F | p | VS-MPR |
| **R1 (L. PFrdls/PFcdl)** | 1.882 | 0.173 | 1.212 | 3.709 | 0.058 | 2.234 | 1.925 | 0.169 | 1.224 | 0.381 | 0.538 | 1.000 |
| **R2 (L. PFcdl)** | 0.182 | 0.670 | 1.000 | 2.840 | 0.095 | 1.641 | 0.226 | 0.635 | 1.000 | 0.420 | 0.519 | 1.000 |
| **R3 (L. PFrdls)** | 1.602 | 0.209 | 1.124 | 1.999 | 0.161 | 1.250 | 0.398 | 0.530 | 1.000 | 1.008 | 0.321 | 1.009 |
| **R4 (L. PFrd)** | 0.200 | 0.656 | 1.000 | 1.523 | 0.220 | 1.104 | 5.998 | 0.016 | 5.455 | 1.102 | 0.297 | 1.020 |
| **R5 (L. PFcdm)** | 0.116 | 0.734 | 1.000 | 6.054 | 0.016 | 5.620 | 1.433 | 0.235 | 1.082 | 4.922 | 0.029 | 3.585 |
| **R6 (R. PFrd)** | 2.368 | 0.127 | 1.402 | 0.542 | 0.463 | 1.000 | 0.029 | 0.865 | 1.000 | 0.192 | 0.662 | 1.000 |
| **R7 (R. PFcdm)** | 0.060 | 0.807 | 1.000 | 2.875 | 0.093 | 1.661 | 0.253 | 0.616 | 1.000 | 1.530 | 0.219 | 1.106 |
| **R8 (R. PFrdls)** | 0.263 | 0.609 | 1.000 | 0.051 | 0.822 | 1.000 | 0.111 | 0.740 | 1.000 | 0.240 | 0.626 | 1.000 |
| **R9 (R. PFcdl)** | 0.188 | 0.666 | 1.000 | 0.0005 | 0.981 | 1.000 | 0.0002 | 0.990 | 1.000 | 2.105 | 0.150 | 1.292 |
| **R10 (R. PFrdls/PFcdl)** | 2.398 | 0.125 | 1.415 | 3.158 | 0.079 | 1.836 | 0.541 | 0.464 | 1.000 | 0.014 | 0.907 | 1.000 |

VS-MPR – Vovk-Sellke maximum p-ratio; O2Hb – Oxygenated Hemoglobin; HHb – Deoxygenated Hemoglobin; R. – Right hemisphere; L. – Left hemisphere; PFrdls – Rostral dorsolateral superior prefrontal cortex; PFcdl – Caudal dorsolateral prefrontal cortex; PFrd – Rostrodorsal prefrontal cortex; PFcdm – Caudal dorsomedial prefrontal cortex
